# The Microglial TREM2 Receptor Programs Hippocampal Development in a Mouse Model of Childhood Deprivation

**DOI:** 10.1101/2025.08.11.669425

**Authors:** Sahabuddin Ahmed, Christian Bowers, Jose Munoz-Martin, Sumit Jamwal, Basavaraju G. Sanganahalli, Lauryn Giuliano, Zoë A. MacDowell Kaswan, Fahmeed Hyder, X. William Yang, Arie Kaffman

## Abstract

Childhood neglect and deprivation are the most common forms of adversity, yet their biological impact on cognitive development—and how enrichment mitigates these effects—remains unclear. Using limited bedding (LB) as a mouse model of deprivation, we previously showed that abnormal microglial-mediated synaptic pruning during the second and third postnatal weeks leads to impaired synaptic connectivity and hippocampal dysfunction, particularly in males.

Here, we demonstrate that LB reduces expression of Triggering Receptor Expressed on Myeloid cells 2 (TREM2) in different mouse strains and that TREM2 deficiency contributes to, but does not fully explain, impaired microglial pruning. Overexpressing TREM2 restored microglial phagocytic function and rescued deficits in hippocampal connectivity and fear learning. Brief postnatal enrichment (P14–P17) also normalized synaptic pruning in a TREM2-dependent manner. Together, our findings identify TREM2 as a key molecular mediator of experience-dependent plasticity, revealing its central role in linking early-life deprivation and enrichment to cognitive outcomes later in life.

## Introduction

Early-life adversity (ELA) encompasses a heterogeneous set of childhood hardships, including abuse, neglect, severe poverty, and exposure to high levels of neighborhood crime^1–6^. ELA is one of the most significant and preventable contributors to abnormal brain development and psychopathology later in life^4,7^, accounting for nearly half of all childhood psychiatric diagnoses^8^ and an estimated annual cost of $2 trillion in the United States^9^. Among the 546,159 cases documented by the Department of Health and Human Services in 2023, 69% were attributed to neglect and deprivation, compared with 10.5% for physical abuse and 7.5% for sexual abuse^10^. Neglect and deprivation are associated with distinct structural and functional brain abnormalities not typically observed in other forms of adversity, including stunted growth, cortical thinning, hyperactivity, and profound cognitive deficits^11–15^—particularly impairments in hippocampus-dependent episodic memory^16,17^. Although early interventions such as adoption and enrichment programs can improve some outcomes^11,18–20^, the cellular and molecular mechanisms by which deprivation and enrichment affect brain development and cognition remain poorly understood.

Despite being the most prevalent form of ELA, neglect is the least studied^11,15^, and no existing animal models have fully captured the key features of childhood deprivation or neglect^21,22^. Furthermore, existing enrichment research has primarily focused on adult animals^23^, with only a handful of studies investigating the effects of early enrichment on cognitive and structural deficits induced by ELA^24–27^. To address these gaps, we recently demonstrated that mice raised under sustained impoverished conditions of limited bedding (LB) exhibit outcomes resembling those seen in children exposed to neglect, including stunted growth, abnormal attachment, cortical thinning, hyperactivity, and deficits in hippocampal-dependent function^21,28,29^. Although most abnormalities were observed in both sexes^21,28,29^, deficits in synaptic connectivity and hippocampal function were significantly more pronounced in LB males compared to LB female littermates^29^. These included a reduced synaptic maturity index (the ratio of mature to immature spines), lower glutamatergic synapse density, and decreased local functional connectivity measured using resting-state fMRI^29^. The sex differences in hippocampal function observed in our studies align with by previous work from other laboratories and by neuroimaging data in humans^1,30,31^.

The high number of immature spines suggested that LB impairs microglia-mediated synaptic pruning^29,32^. We found significant pruning deficits in postnatal day 17 (P17) pups, when pruning peaks in the hippocampus^29,33–37^. Transient microglial elimination during the second and third weeks of life reproduced the sex-specific cognitive and synaptic abnormalities observed in adolescent LB mice, whereas chemogenetic activation of microglia during the same period normalized pruning at P17 and rescued structural, functional, and behavioral deficits^29^. Although LB impaired microglial pruning in both sexes, LB females had enhanced astrocyte-mediated pruning, providing a potential compensatory mechanism against microglial dysfunction^29^. These findings indicate that disrupted microglia-mediated synaptic pruning during the second and third weeks of life directly underlies the synaptic connectivity and hippocampal functional deficits seen in adolescent LB males^32^.

Microglia isolated from the developing hippocampus of LB mice exhibited reduced expression of the TREM2 receptor^38^. This is significant because TREM2 is exclusively expressed in microglia and is essential for normal synaptic pruning in the developing hippocampus^35,36,38^, as well as for maintaining synaptic connectivity and social behavior later in life^35^. However, the precise role of microglial TREM2 in mediating the synaptic and behavioral deficits observed in LB male mice has not been fully elucidated. Here, we show that LB impairs microglial synaptic pruning through both TREM2-dependent and TREM2-independent mechanisms. Nevertheless, overexpression of TREM2 in microglia was sufficient to normalize synaptic pruning during this critical developmental period and to rescue deficits in synaptic connectivity and hippocampal-dependent behavior later in life.

Since LB appears to model childhood deprivation, we tested whether adding toys to the cages of LB mice could increase TREM2 expression and normalize microglia-mediated synaptic pruning in the developing hippocampus. Indeed, we found that postnatal enrichment from P14 to P17 normalized TREM2 expression and phagocytic activity in LB Trem2-wildtype mice, but not in LB Trem2-knockout mice. Together, these findings reveal a novel role for TREM2 in mediating the synaptic and cognitive consequences of early-life deprivation, as well as the restorative effects of early enrichment on microglia-mediated synaptic pruning during a critical period of hippocampal development

## Results

### LB Reduces Microglial TREM2 Expression and Impairs Microglia-Mediated Synaptic Pruning in the Developing Hippocampus Across Multiple Mouse Strains

Exposure to LB impairs the ability of microglia to phagocytose synaptic material during the critical period of hippocampal synaptic pruning, leading to deficits in synaptic connectivity and hippocampal function later in life^29,38^. Specifically, LB reduced microglial cell volume, phagosome size, and the number of PSD95-positive synaptic puncta engulfed by microglia in the hippocampus of P17 pups, a time point when synaptic pruning peaks in this region^29,33–37^. This abnormal phagocytic activity was accompanied by reduced expression of the TREM2 receptor, which is necessary for normal synaptic pruning during hippocampal development^29,38^. Since these findings were initially obtained in BALB/c mice, we sought to determine whether they could be replicated in C57BL/6J (C57) mice, a strain commonly used for transgenic studies. Here, we show that C57 P17 male and female pups exposed to LB exhibit similar deficits in microglia-mediated synaptic pruning as previously observed in BALB/c mice (Fig. 1A-F). Furthermore, using a TREM2-specific antibody, we confirmed that TREM2 expression is reduced in microglia localized to the stratum radiatum of 17-day-old C57 LB pups (Fig. 1 G-I). Together, these findings demonstrate that LB impairs microglia-mediated synaptic pruning and reduces TREM2 expression during a critical period of hippocampal development across multiple mouse strains.

**Figure 1.**
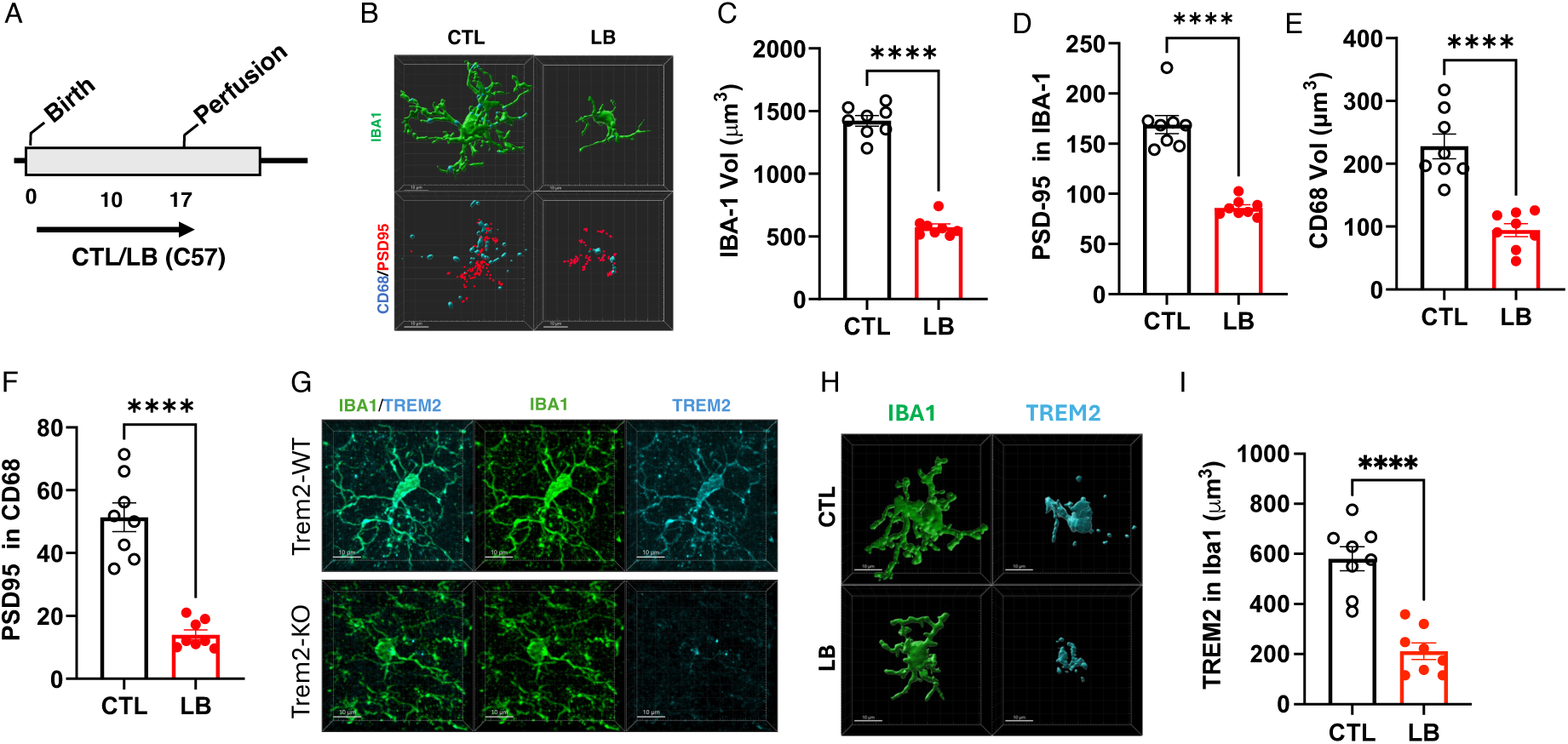
Phagocytic activity and TREM2 expression in P17 C57BL/6J pups. (**A**) Experimental timeline. (**B**) Representative Imaris reconstruction showing Iba1 (green), CD68 (cyan), and PSD95 puncta (red) staining of microglia in the stratum radiatum of P17 mice. (**C**) Iba1 volume, t(14) = 16.80, P < 0.0001. (**D**) PSD95-positive puncta within Iba1+ cells, t(14) = 8.71, P < 0.0001. (**E**) CD68 volume (phagosome), t(14) = 6.06, P < 0.0001. (**F**) PSD95 puncta within CD68+ phagosomes, t(14) = 7.76, P < 0.0001. (**G**) Representative images of Iba1 (green) and TREM2 (cyan) staining in Trem2-WT (top) and Trem2-KO (bottom) P17 mice raised under CTL conditions. **(H**) Representative Imaris reconstruction images of Iba1 (green) and TREM2 (cyan) staining in CTL and LB mice. (**I**) Quantification of TREM2 expression levels in microglia from CTL and LB mice. Data are presented as mean ± SEM and analyzed using Student’s *t*-tests, P > 0.05 (not shown), * P < 0.05, ** P < 0.01, *** P < 0.001, **** P < 0.0005. *N* = 8 mice per rearing condition (sex-balanced) from 4 independent litters.

### TREM2 Mediates Some, but Not All, of the Phagocytic Deficits Observed in LB Mice

Work from our group and others has shown that TREM2 is necessary for normal microglia-mediated synaptic pruning in developing mouse pups^35,36,38^. However, the contribution of this receptor to the phagocytic deficits observed in microglia from LB mice has not been fully elucidated. To address this question, we exposed *Trem2* wild-type (*Trem2-WT*) and *Trem2* knockout (*Trem2-KO*) mice to CTL and LB rearing conditions. At P17, all pups were perfused to assess the effects of rearing, genotype, and their interaction on microglial cell volume, phagosome size, and the number of PSD95-positive puncta within microglia (Fig. 2A). Consistent with previous work^29,38^, there were no significant main effects or interactions involving sex, so data from both sexes were combined and analyzed using a two-way ANOVA examining the effects of rearing, genotype, and their interaction. A significant interaction was found between rearing and genotype (F(1, 28) = 48.80, P < 0.0001). A Tukey-HSD post hoc analysis revealed a 60% total reduction in microglial volume between CTL-*Trem2-WT* and LB-*Trem2-WT* mice. This reduction was attributable to a 30% decrease between CTL-*Trem2-WT* and CTL-*Trem2-KO* mice and an additional 30% decrease between CTL-*Trem2-KO* mice and LB-*Trem2-KO* mice (Fig. 2B-C). Similar outcomes were observed for the number of PSD95 puncta within microglia (Fig. 2B-D). These findings replicate our previous observations of reduced microglial volume and phagocytic activity in *Trem2-KO* mice under CTL conditions^38^ and suggest that lower TREM2 levels account for approximately half of the reduction in cell volume and phagocytic activity observed in LB mice (i.e., a TREM2-dependent mechanism). Notably, in the absence of TREM2, LB exposure led to an additional 30% reduction in microglial volume, suggesting the involvement of a TREM2-independent mechanism in microglia-mediated synaptic pruning (Fig. 2B-D). In contrast, reduction in TREM2 expression (TREM2-dependent mechanism) accounted for most of the effects of LB on phagosome size and the number of PSD95 puncta within the phagosome, with minimal impact of LB in Trem2-KO mice on these measures (Fig. 2E-F). In summary, reduced TREM2 expression accounts for approximately half of the deficits in synaptic material clearance and most of the deficits in phagosome size and function observed in LB mice.

**Fig 2.**
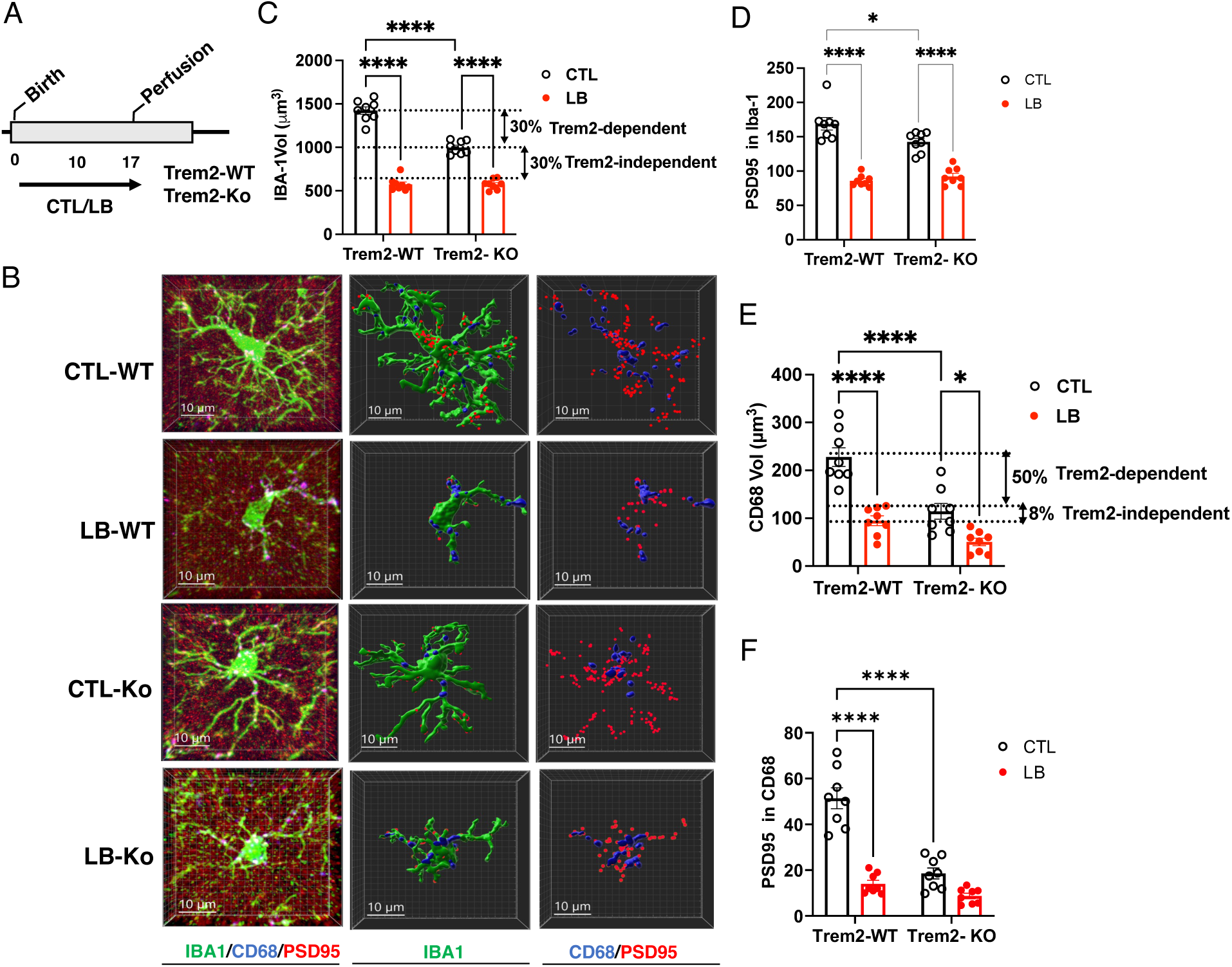
Trem2 Mediates Some of the Phagocytic Deficits Observed in LB mice. (**A**) Experimental Timeline. (**B**) Representative confocal images and Imaris reconstructions of microglia stained with IBA1 (green), CD68 (blue), and PSD95 puncta (red) located in the stratum radiatum of P17 mice from *Trem2-WT* (WT) and *Trem2-KO* (Ko) animals exposed to either control (CTL) or LB conditions. (**C**) Microglial volume. Interaction: F (1, 28) = 7.719, P = 0.0096. (**D**) PSD95 Puncta engulfed by microglia. Interaction: F (1, 28) = 7.72, P = 0.0096. (**E**) Phagosome CD68 volume. Interaction: F (1, 28) = 5.88, P = 0.022. (**F**) PSD95 puncta inside CD68. Interaction: F (1, 28) = 24.88, P < 0.0001. Data are presented as mean ± SEM and analyzed using 2-way ANOVA. Tukey-HSD post-hoc: P > 0.05 (not shown), * P < 0.05, ** P < 0.01, *** P < 0.001, **** P < 0.0005. N=8 mice from 6 litters per rearing and genotype group, half of which are females.

### TREM2 Overexpression Restores Normal Phagocytic Activity in P17 LB Mice

Overexpressing human TREM2 in microglia has been found to restore normal phagocytic activity in aged Alzheimer’s mouse models, enabling more efficient clearance of amyloid beta deposits and improved cognition^39^. Using these mice, we tested whether overexpression of TREM2 could rescue the microglia-mediated synaptic pruning deficits observed in the developing hippocampus of LB mice. This was achieved by mating heterozygous *Trem2-OE* males with *Trem2* wildtype females (*Trem2-WT*) to generate mixed litters containing *Trem2-WT* and *Trem2-OE* offspring. Litters were then randomly assigned to either CTL or LB conditions and perfused at P17 to assess the effects of rearing, genotype, and sex on TREM2 expression and in vivo phagocytic activity. No significant main effects or interactions with sex were found for any dependent variable; thus, data from male and female mice were combined. Initial analyses revealed a significant interaction between rearing and genotype for microglial volume, and TREM2 expression, driven by reduced microglial volume and TREM2 expression in LB-*Trem2-WT*, but not LB-*Trem2-OE* littermates (Fig. S1 A-C). Notably, microglial volume and TREM2 levels were comparable among CTL-*Trem2-WT*, CTL-*Trem2-OE*, and LB-*Trem2-OE* mice, indicating that this manipulation normalized microglial volume and TREM2 expression in P17 LB mice without affecting these measures in CTL mice (Fig. S1 A-C). TREM2 overexpression also normalized the stunted growth observed in LB mice, consistent with previous findings that chemogenetic activation of microglia increases body weight in P17 LB pups^29^ (Fig. S1D). Similar results were obtained in a second cohort tested for in vivo phagocytic activity, with *Trem2-OE* mice able to restore normal phagocytic activity in LB mice (Fig. 3). Thus, overexpression of TREM2 in microglia is sufficient to correct microglia-mediated synaptic deficits during a critical period of hippocampal development in mice raised under impoverished LB conditions.

**Fig. 3.**
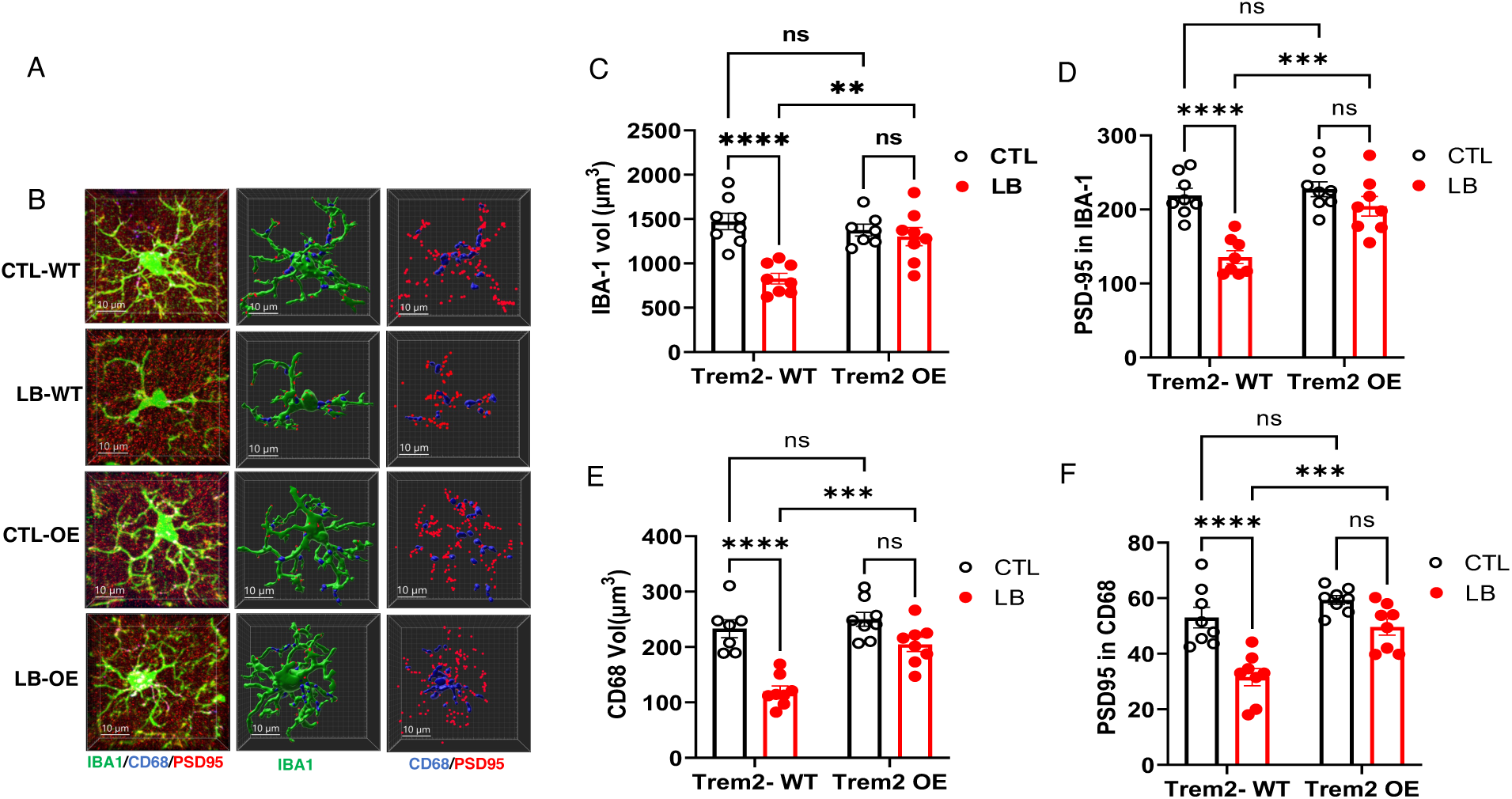
Overexpression of TREM2 Normalize Microglial-Mediated Synaptic Pruning in the Developing Hippocampus of LB Mice. (**A**) Experimental Timeline. (**B**) Representative confocal and Imaris images of microglia stained with IBA1 (green), CD68 (blue), and PSD95 puncta (red) located in the stratum radiatum of P17 mice from *Trem2-WT* (WT) and *Trem2-OE* (OE) mice exposed to control (CTL) or LB conditions. (**C**) Microglial volume. Rearing x genotype interaction: F (1, 27) = 11.60, P = 0.0021. (**D**) PSD95 Puncta engulfed by microglia. Interaction: F (1, 28) = 8.07, P = 0.0083. (**E**) Phagosome CD68 volume. Interaction: F (1, 27) = 6.91 = 1.28, P = 0.014. (**F**) PSD95 puncta inside CD68. Rearing: F (1, 28) = 28.28, P < 0.0001, Genotype: F (1, 28) = 17.25, P = 0.0003, Interaction: F (1, 28) = 3.95, P = 0.056. Data are presented as mean ± SEM and analyzed using 2-way ANOVA. NS = non-significant, Post-hoc: P > 0.05 (NS), * P < 0.05, ** P < 0.01, *** P < 0.001, **** P < 0.0005. N = 8 mice from 6 litters, per rearing and genotype group, half of which are females.

### Overexpression of TREM2 Corrects Deficits in Contextual Freezing in LB Males

Given the critical role of microglia-mediated synaptic pruning in programming hippocampal function, we predicted that overexpression of TREM2 would also normalize contextual freezing behavior in adolescent LB male mice. To test this, we randomized mixed *Trem2-WT* and *Trem2-OE* litters to CTL and LB conditions and assessed contextual fear conditioning in P30–33 adolescent mice (Fig. 4A). All mice exhibited a significant increase in freezing behavior during the first day of training, with no significant effects of rearing, sex, or genotype (Fig. S2). A three-way ANOVA for freezing behavior during the second day of contextual testing revealed a significant rearing x sex interaction (F(1, 97) = 3.88, P = 0.05), prompting separate analyses for males and females. In males, a two-way ANOVA revealed a significant interaction between rearing and genotype, driven by a reduction in freezing behavior in LB-*Trem2-WT* mice compared to all other groups. Freezing behavior was comparable among CTL-*Trem2-WT*, CTL-*Trem2-OE*, and LB-*Trem2-OE* groups (Fig. 4B), indicating that *Trem2-OE* was able to normalize this hippocampal-dependent behavior in adolescent LB males. No significant main effects or interactions were detected in females (Fig. 4C), a finding consistent with our previous work^21,29^.

**Fig. 4.**
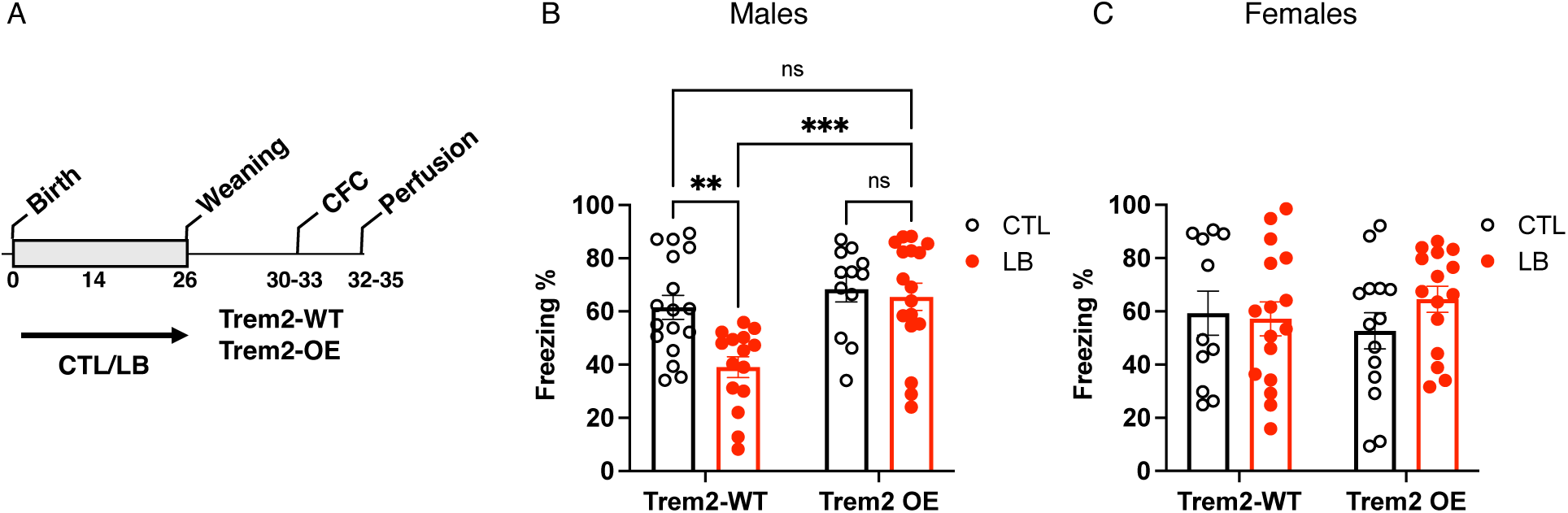
Trem2 OE Normalizes Contextual Freezing in Males. **(A)** Timeline **(B)** CFC males. Rearing x genotype interaction (1, 57) = 4.350, P = 0.041. **(C)** CFC females. Rearing: F (1, 52) = 0.54, P = 0.46, Genotype: F (1, 52) = 0.0027, P = 0.96, Interaction: F (1, 52) = 1.134, P = 0.29. Data are presented as mean ± SEM and analyzed using 2-way ANOVA. NS = non-significant, Post-hoc: P > 0.05 (NS), * P < 0.05, ** P < 0.01, *** P < 0.001, **** P < 0.0005. CTL-*Trem2-WT*: males = 12, females = 12, 6-7 litters, CTL-*Trem2-OE*: males = 10, females = 13, 6-7 litters, LB-*Trem2-WT*: males = 13, females = 15, 7-8 litters, LB-*Trem2-OE*: males = 16, females = 14, 7-8 litters.

### Overexpression of TREM2 Mitigates Deficits in Dendritic Varicosity, Synaptic Maturity, and Glutamatergic Synapse Density in Adolescent LB Male Mice

Next, we used DiOlistic labeling to test whether overexpression of TREM2 could also correct the structural synaptic deficits previously observed in adolescent LB male mice^29^ (representative DiOlistic labeled images and spine reconstruction shown in Fig. 5A). Careful assessment of dendritic diameter in CA1 pyramidal neurons within the stratum radiatum revealed a significant increase in diameter in LB-*Trem2-WT* mice compared to all other groups (Fig. 5B). This increase in average diameter was due to abnormal ballooning or varicosities along the dendritic shaft in LB-*Trem2-WT* mice (Fig. S3), which were restored to normal levels in LB-*Trem2-OE* mice (Fig. 5B). Analysis of total spine density revealed a significant main effect of rearing condition, with no significant effect of genotype or interaction (Fig. 5C). A similar pattern was observed for the density of immature spines; however, the trending effect of rearing did not reach significance (P = 0.074; Fig. 5D). The density of mature (mushroom) spines showed significant main effects of both rearing and genotype, but no significant interaction (Fig. 5E). The ratio between mature (mushroom) and immature spines (stubby, thin, filopodia), termed the ’maturity index,’ was reduced in adolescent LB-*Trem2-WT* male mice but not in LB-*Trem2-WT* female mice, a reduction that was directly related to microglial phagocytic activity during the second and third weeks of life^29^. Consistent with these prior findings, we found that TREM2 overexpression fully mitigated the deficits in maturity index observed in LB-*Trem2-WT* males (Fig. 5F).

**Fig 5.**
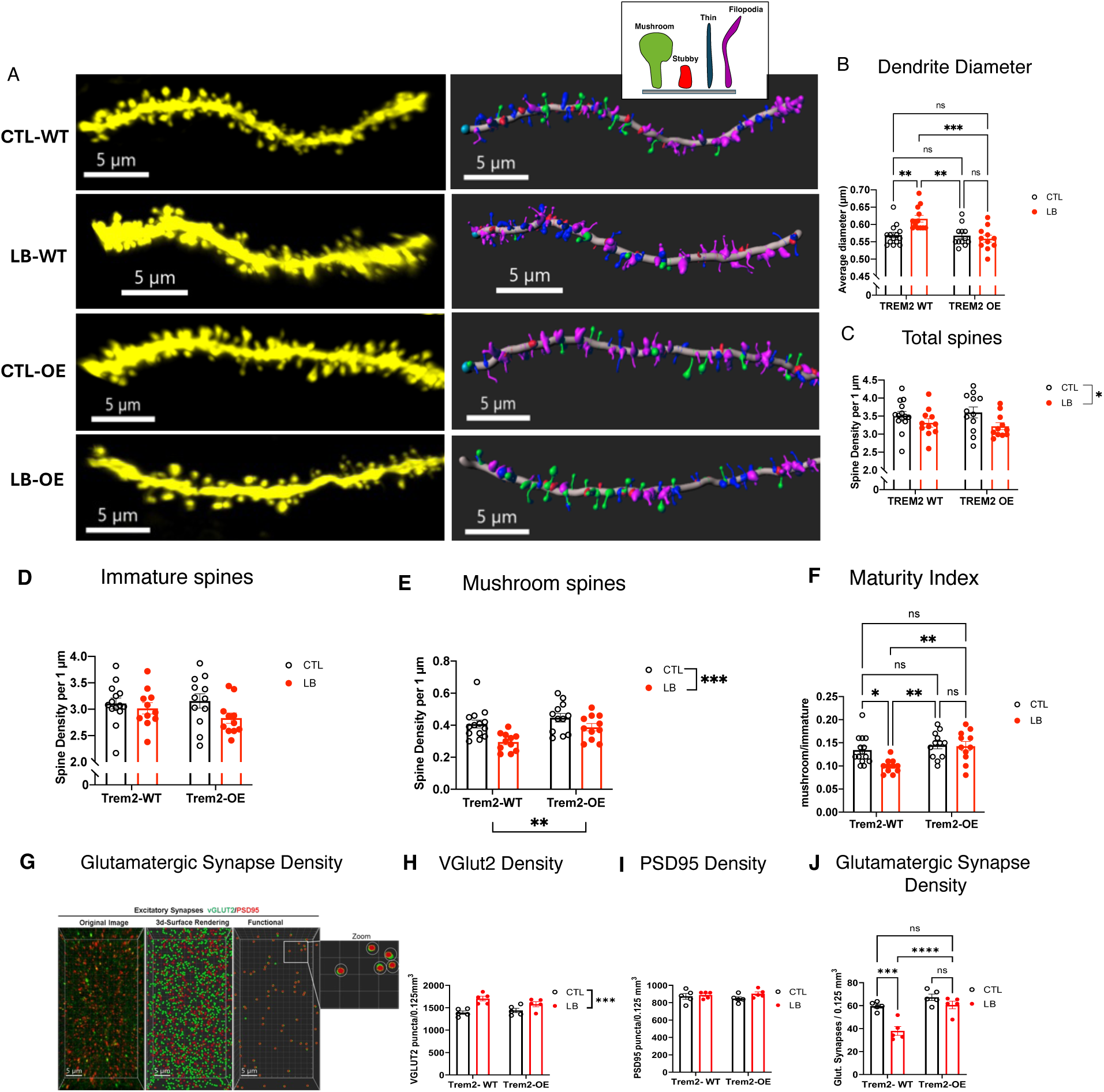
TREM2 Overexpression Restores Deficits in Dendrite Diameter, Synaptic Maturity **and Glutamatergic Synaptic Density in Adolescent LB Males.** (A) Representative confocal images and Imaris models of apical dendrites in the stratum radiatum. Insert: mushroom spines (green), stubby (red), thin spines (blue), filopodia (magenta). (B) Average dendrite diameter. Interaction: F (1, 45) = 9.529, P = 0.0035. (C) Total Spine density. Rearing: F (1, 44) = 5.593, P = 0.022, Genotype: NS, Interaction: NS. (D) Immature Spine density. Rearing: F (1, 44) = 3.35, P = 0.074, Genotype: NS, Interaction: NS. (E) Mushroom Spine density. Rearing: F (1, 44) = 12.66, P = 0.0009, Genotype: F (1, 44) = 7.461, P = 0.0090, Interaction: NS. (F) Maturity Index. Interaction: F (1, 44) = 4.15, P = 0.048. (G) Representative confocal images and Imaris models of VGlut2 and PSD95 puncta in the stratum radiatum. (H) VGlut2 density. Rearing: F (1, 16) = 21.72, P = 0.0003, Genotype: NS, Interaction: F (1, 16) = 3.26, P = 0.089 (I) PSD95 density. Rearing: NS, Genotype: NS, Interaction: NS (J) Glutamatergic synapse density. Rearing: Interaction: F (1, 16) = 6.061, P = 0.025. Data are mean ± SEM and analyzed using 2-way

Next, we tested whether TREM2 overexpression could also ameliorate deficits in glutamatergic synapse density in the stratum radiatum of LB male mice (Fig. 5G–J). Two-way ANOVA revealed a significant main effect of rearing, but no significant effect of genotype or interaction, on the density of the presynaptic marker VGlut2 (Fig. 5H). No significant effects of rearing, genotype, or interaction were found for the postsynaptic marker PSD95 (Fig. 5I). However, the density of putative functional glutamatergic synapses—defined as colocalized pre-and postsynaptic markers—was reduced in LB-*Trem2-WT* mice and was restored to levels comparable to CTL-*Trem2-WT* in LB-*Trem2-OE* mice (Fig. 5J). Hence, microglial TREM2 overexpression was sufficient to normalize the increase in dendritic varicosity, the reduction in maturity index, and the decreased glutamatergic synapse density observed in the stratum radiatum of adolescent LB male mice.

### TREM2 Overexpression Restores Local Functional Connectivity in the Right Dorsal Hippocampus of LB Male Mice

Using whole-brain rsfMRI voxel analysis to unbiasedly identify brain regions with altered local functional connectivity, we recently found reduced local connectivity in multiple brain regions—including the dorsal hippocampus—in adolescent LB male mice, but not in LB female littermates^29^. We therefore tested whether TREM2 overexpression could rescue these connectivity deficits in the dorsal hippocampus. To do this, we examined differences in local functional connectivity across rearing conditions in *Trem2-WT* males (CTL-WT vs. LB-WT) and *Trem2-OE* males (CTL-OE vs. LB-OE), applying rigorous correction for multiple comparisons (see Statistical Analysis and previous work). Consistent with earlier findings, LB reduced local functional connectivity in the dorsal hippocampus of *Trem2-WT* mice (Fig. 6A) but not in *Trem2-OE* mice (Fig. 6B). Interestingly, these deficits in *Trem2-WT* mice were primarily observed in the right dorsal hippocampus (Fig. 6C-D).

**Fig 6.**
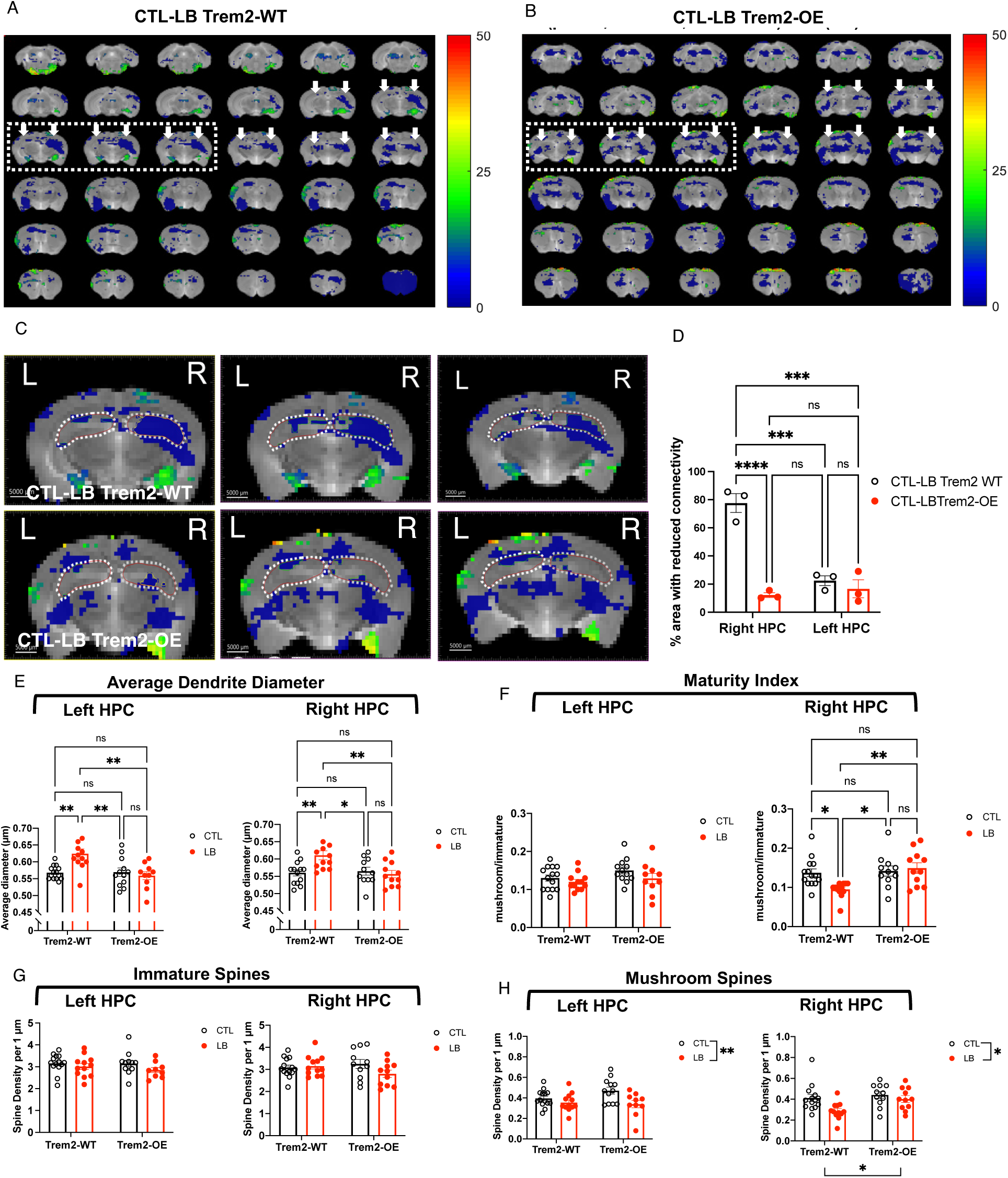
TREM2 Overexpression Normalizes Right Dorsal Hippocampal Connectivity in Adolescent LB Male Mice. Effects of rearing on local functional connectivity in *Trem2-WT* males (**A**) and *Trem2-OE* males **(B**). Blue and green colors represent brain regions with significant reduction in local functional connectivity P < 0.05, FDR corrected, local cluster size k> 25 voxels. White arrows indicate the location of the hippocampus (HPC), and the dotted areas are the slices used for further analysis in C-D. (**C**) The percentage of reduced local functional connectivity area (in blue) within the left and right HPC (dotted lines) were calculated using Imaris and are shown in (**D**). (**D**) Percent area of reduced local functional connectivity, genotype x side interaction: F (1, 8) = 35.35, P = 0.0003. right hpc-*Trem2-WT* vs. all other groups (left hpc-*Trem2-WT*, right hpc-*Trem2-OE*, left hpc-*Trem2-OE*) P < 0.001, no significant differences between left hpc-*Trem2-WT*, right hpc-*Trem2-OE*, and left hpc-*Trem2-OE*. (E-H) DiOlistic labeling. (**E**) Average Dendritic Diameter. Left HPC: rearing x genotype Interaction: F (1, 44) = 8.81, P = 0.0048. Right HPC: Interaction: F (1, 43) = 6.991, P = 0.011. (**F**) Maturity Index. Left HPC: Rearing: NS, Genotype: NS, Interaction: NS. Right HPC: Interaction: F (1, 44) = 5.28, P = 0.026. (**G**) Immature Spine Density. Left HPC: Rearing: F (1, 44) = 4.20, P = 0.046, Genotype: NS, Interaction: NS. Right HPC: Rearing: NS, Genotype: NS, Interaction: F (1, 44) = 2.86, P = 0.098. (E) Mushroom Spine Density. Left HPC: Rearing: F (1, 44) = 8.60, P = 0.0053, Genotype: NS, Interaction: NS. Right HPC: Rearing: F (1, 44) = 6.685, P = 0.013, Genotype: F (1, 44) = 5.19, P = 0.027, Interaction: NS. Data are mean ± SEM and analyzed using 2-way ANOVA. Post-hoc: P > 0.05 (non-significant, NS), * P < 0.05, ** P < 0.01, *** P < 0.001, **** P < 0.0005. N = 6 males per rearing and genotype group (A-D). N = 11-14 male mice per rearing and genotype group (E-H).

Moreover, using an independent cohort of mice (the same cohort used in Fig. 5 for DiOlistic labeling), we re-examined the effects of rearing and genotype in the left and right dorsal hippocampus. A similar pattern was found for average dendritic diameter in both hemispheres, confirming that TREM2 overexpression rescued varicosity deficits bilaterally (Fig. 6E). In contrast, LB-mediated deficits in the maturity index were observed only in the right hippocampus of *Trem2-WT* mice and were corrected in *Trem2-OE* mice (Fig. 6F), supporting the lateralization found using rsfMRI. No significant effects of rearing, genotype, or their interaction were found for the density of immature synapses in either hemisphere (Fig. 6G). For the density of mushroom spines, a two-way ANOVA revealed significant main effect of rearing, but no significant effect of genotype or interaction in the left hippocampus; significant effects of rearing and genotype, but no interaction was found in the right hippocampus (Fig. 6H). Finally, no significant differences in local functional connectivity were found when a similar analysis was conducted in female mice (Fig. S4), confirming our previous findings^29^.

### Postnatal Enrichment Restores Synaptic Pruning via TREM2-Dependent Microglial Mechanisms

Given that LB induces structural and behavioral deficits reminiscent of those observed in children and adolescents exposed to severe neglect and deprivation^21,22^, we hypothesized that environmental enrichment during this critical period of hippocampal development could mitigate these deficits. To test this, we introduced toys into a subset of LB cages at P14—a condition termed LB + Toys (LBT)—and perfused the animals at P17 to assess microglia-mediated synaptic pruning in the stratum radiatum. Initial testing in BALB/c mice indicated that three days of enrichment fully normalized the number of PSD95 puncta engulfed by microglia (Fig. S5A-E) and in the phagosome (Fig. S5F), despite only partially correcting deficits in total cell volume (Fig. S5G) and having no significant impact on phagosome size (Fig. S5H). We then repeated these studies in *Trem2-WT* and *Trem2-KO* mice (C57 background, Fig 7A). LBT in *Trem2-WT* mice was sufficient to normalize microglial TREM2 expression (Fig. 7B-C) and phagocytic activity (Fig. 7D-H) at P17. Specifically, the number of PSD95 puncta engulfed by microglia in LBT-*Trem2-WT* mice was significantly greater compared to LB-*Trem2-WT* mice, reaching levels comparable to CTL-*Trem2-WT* mice (Fig. 7E), with similar outcomes observed for PSD95 puncta within the phagosome (Fig. 7F). Enrichment partly rescued total microglial volume (Fig. 7G) and had no significant effect on phagosome size (Fig. 7H). LBT had no effect on any of these variables in Trem2-KO mice (C57 background), indicating that upregulation of TREM2 is necessary for the therapeutic effect of postnatal enrichment on microglia-mediated synaptic pruning.

**Fig. 7.**
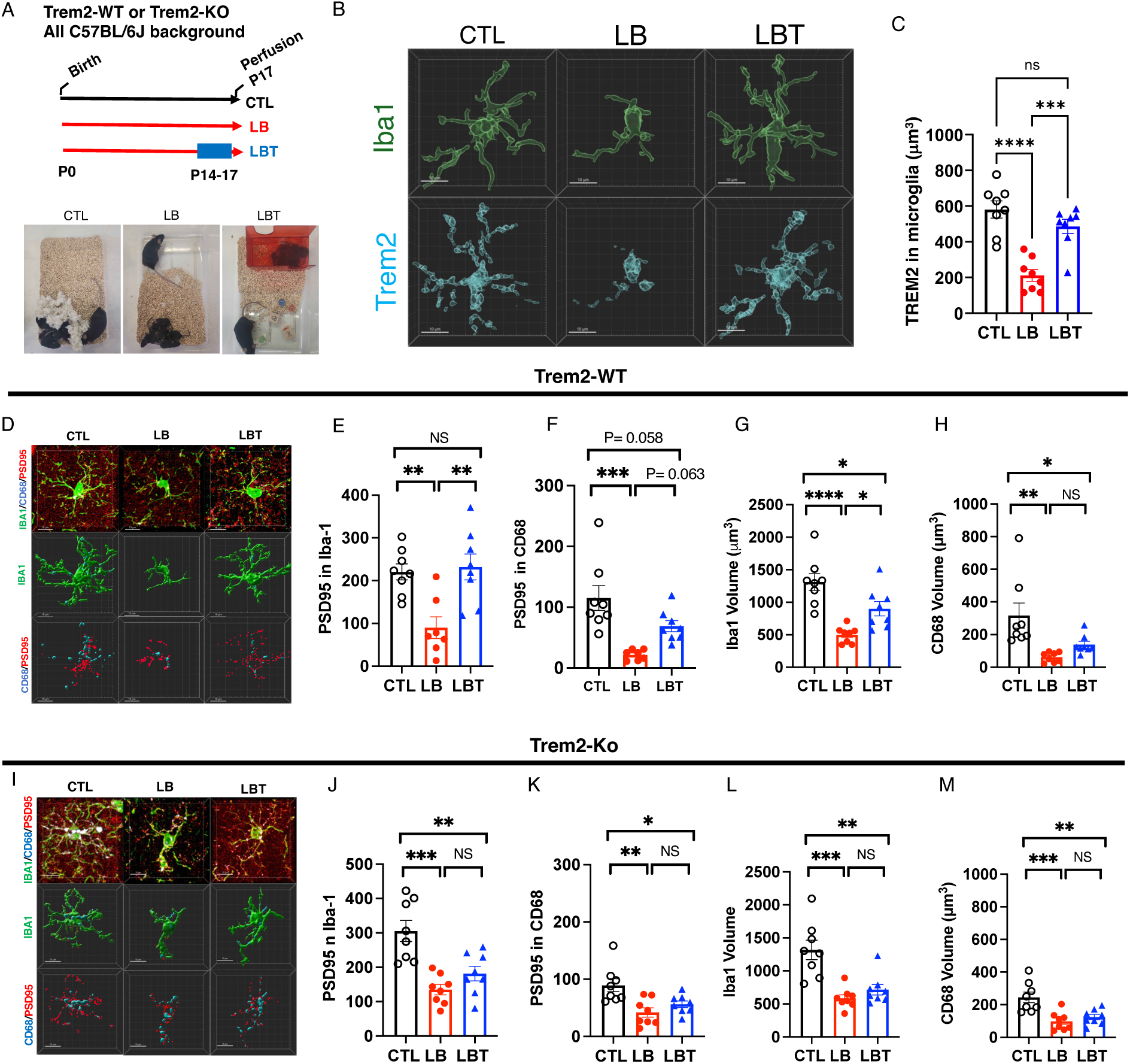
TREM2 Is Essential for Enrichment-Induced Restoration of Microglial Function and Morphology. **(A**) Experimental timeline and images of CTL, LB and LBT rearing conditions. **(B)** Representative Imaris images of IBA1 (green) and TREM2 (cyan) expression in CTL, LB and LBT, P17 *Trem2-WT* pups**. (C)** TREM2 expression in microglia, Rearing: F (2, 21) = 21.98, P < 0.0001. **(D-H)** Effects of rearing in microglial-mediated synaptic pruning in Trem2-WT P17 pups. (**D**) Representative Imaris reconstructions of microglia stained with IBA1 (green), PSD95 puncta (red) and CD68 (cyan) taken from stratum radiatum of *Trem2-WT* mice. (**E**) PSD95 puncta in microglia-*Trem2-WT*: Rearing F (2, 20) = 9.249, P = 0.0014. (**F**) PSD95 puncta in phagosome-*Trem2-WT*: Rearing F (2, 20) = 11.47, P = 0.0005. (**G**) Microglial volume-*Trem2-WT*: Rearing F (2, 21) = 15.91, P < 0.0001. (**H**) Phagosome volume-*Trem2-WT*: Rearing F (2, 21) = 7.99, P = 0.0026. **(I-M)** Effects of rearing in microglial-mediated synaptic pruning in *Trem2-KO* P17 pups. (**I**) Representative Imaris reconstructions of microglia stained with IBA1 (green), PSD95 puncta (red) and CD68 (cyan) taken from stratum radiatum of *Trem2-KO* mice. **(J)** PSD95 puncta in microglia-*Trem2-KO*. Rearing F (2, 21) = 14.23, P=0.0001. **(K)** PSD95 puncta in phagosome-*Trem2-KO*. Rearing F (2, 21) = 7.74, P = 0.0030. **(L)** Microglial volume-*Trem2-KO*. Rearing F (2, 21) = 14.75, P < 0.0001. **(M)** Phagosome volume-*Trem2-KO*. Rearing: F (2, 21) = 10.80, P = 0.0006. Data are mean ± SEM and analyzed using a one-way ANOVA. Post-hoc: P > 0.05 (non-significant, NS), * P < 0.05, ** P < 0.01, *** P < 0.001, **** P < 0.0005. N = 8 mice per rearing and genotype, half females.

## Discussion

Despite being the most prevalent form of childhood adversity little is currently known about the underlying biology by which childhood neglect and deprivation impair normal neurodevelopment and cognition or how early interventions can mitigate these deficits^11,15,21,22^. This gap in our knowledge primarily stems from the absence of animal models that can faithfully replicate core features of childhood neglect and deprivation sequalae, while demonstrating reversal through early enrichment^21,22^.

In response to these challenges, we recently found that mice reared under prolonged impoverished conditions using limited bedding (LB) display behavioral and neurobiological outcomes similar to those observed in children exposed to severe neglect, including reduced body weight, insecure attachment behaviors, heightened anxiety, cortical thinning, hyperactivity, and impairments in hippocampal-dependent functions^21,28,38^. Although most abnormalities were observed in males and females, deficits in synaptic connectivity and hippocampal function were more pronounced in adolescent LB males^29^. These sex-specific effects on hippocampal function are consistent with work from other groups and imaging data in humans^1,30,31^. Further investigation revealed that the synaptic and cognitive deficits observed in adolescent LB males were due to abnormal microglia-mediated synaptic pruning during the second and third weeks of life, when hippocampal pruning peaks^29^. LB disrupted microglial synaptic pruning in both sexes; however, LB females appeared to compensate by upregulating astrocyte-mediated pruning, potentially mitigating the effects of microglial dysfunction^29^.

Here we demonstrate that reduced expression of the receptor TREM2 plays a key role in the abnormal microglial-mediated synaptic pruning observed during this critical period of hippocampal development. Overexpression of TREM2 in microglia was sufficient to restore microglial phagocytic function and correct the stunted growth observed in P17 LB mice.

Furthermore, TREM2 overexpression normalized synaptic connectivity and contextual fear conditioning deficits in adolescent LB males. Notably, brief enrichment from P14 to P17 also rescued microglial-mediated synaptic pruning, an effect that required upregulation of TREM2 expression.

Together, these studies suggest a novel conceptual framework for understanding the impact of childhood deprivation and enrichment on long-term changes in connectivity and cognition (Fig 8). Specifically, we propose that high levels of microglial-mediated synaptic pruning during the second and third weeks of life are necessary to remove supernumerary poorly functioning synapses on apical dendrites of CA1 neurons, a process that requires high levels of TREM2 expression by microglia. Inefficient pruning during this critical period, such as occurs under LB conditions, reduces the synaptic maturity index, glutamatergic synaptic density, and local functional connectivity, and impairs contextual fear conditioning later in life. The mechanisms by which synaptic pruning enhances connectivity and hippocampal function is yet to be clarified, but we have previously proposed that the removal of immature spines redirects limited neuronal/apical resources such as Calcium/Calmodulin-dependent protein kinase II (CamKII) or mitochondria towards more functional spines leading to increases in the maturity index and connectivity, and improved hippocampal-dependent function (Fig 8)^32^.

**Fig. 8.**
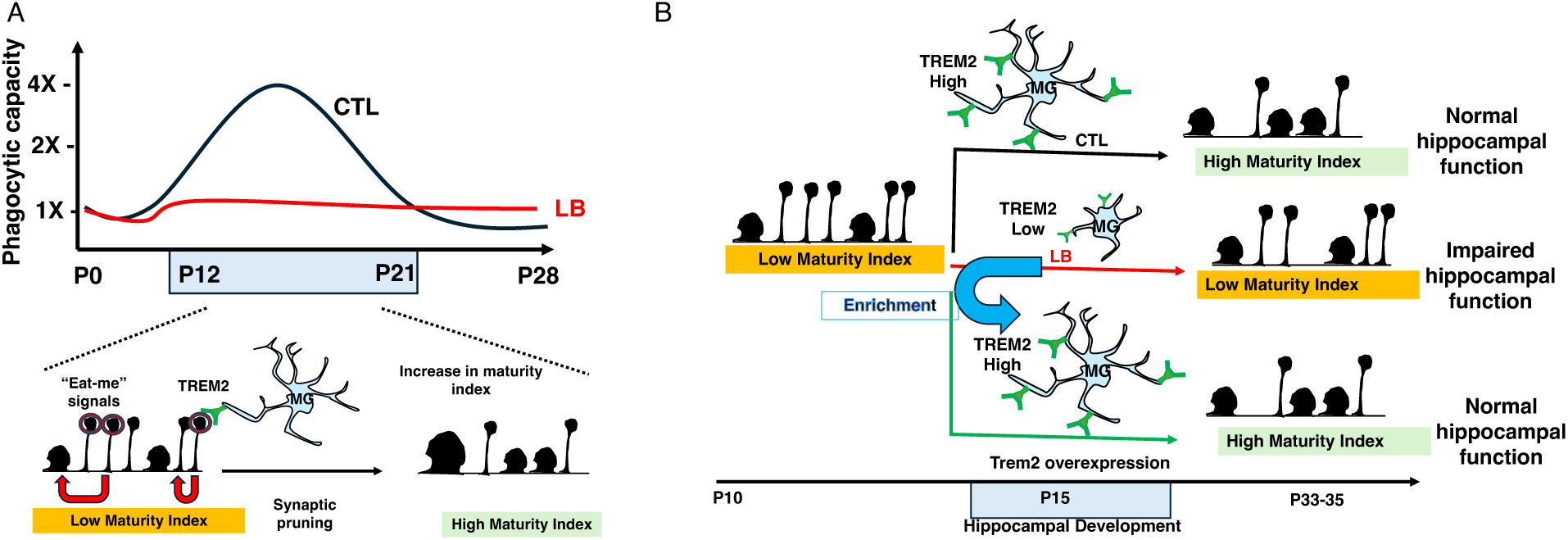
Working Model. **(A)** Top: Microglial capacity to eliminate surplus synapses in the normally developing hippocampus peaks during the second and third weeks of life. This process requires adequate sensory and cognitive activation of CA1 pyramidal neurons, which does not occur in the impoverished rearing conditions experienced by LB pups. Bottom: During this critical period, TREM2 expression on microglia increases, enabling efficient removal of non-functional synapses labeled with “eat-me” signals (red circles). This facilitates the redirection of neuronal resources toward functional synapses (red arrows), resulting in a higher synaptic maturity index and improved hippocampal-dependent memory. **(B)** Reduced TREM2 expression accounts for some, but not all, of the phagocytic deficits observed in LB mice. TREM2 overexpression corrects synaptic and cognitive deficits in adolescent LB mice, while brief enrichment during this critical period normalizes microglial phagocytic activity in a TREM2-dependent manner.

The pathways by which LB inhibits microglial phagocytic activity and reduces TREM2 expression remain unclear. However, the finding that brief enrichment restores phagocytic activity in LB pups in a TREM2-dependent manner suggests that efficient synaptic pruning requires sufficient activation of CA1 neurons during the second and third weeks of life. This level of neuronal activation may not be achieved under the impoverished conditions characteristic of the LB environment. This assertion is supported by previous work demonstrating that exposing LB pups to a novel cage with clean bedding for one hour triggered c-fos activation in CA1 neurons and enhanced microglial ramification and phagocytic activity^38^. Furthermore, high frequency stimulation of the Schaefer collaterals enhances microglial processes outgrowth and synaptic contacts^40^. The close relation between neuronal activation and microglial process extension is mediated by the release of ATP/ADP from activated neurons that in turn recruits microglial process towards active neurons via the P2RY12 purinergic receptor on microglia^41–43^. Chronic whisker trimming early in life--a form of tactile deprivation--impairs synaptic pruning and reduces microglial ramification in the barrel cortex with similar findings obtained with NMDA receptor blockade^44,45^. Together, these findings support the notion that adequate levels of neuronal activation are necessary to support efficient microglial-mediated synaptic pruning. LBT may act to normalize hippocampal microglial function through similar novelty-mediated increases in neuronal activity.

These findings, however, appear inconsistent with studies—primarily in the visual system—showing that sensory deprivation or neuronal inactivity enhances, rather than decreases, microglial-mediated synaptic engulfment^41,46,47^. This deprivation-induced pruning also occurs during critical developmental periods and is thought to serve an adaptive function: eliminating non-functional connections while strengthening functional ones from the unaffected eye or other intact sensory systems. Further research is needed to resolve this apparent discrepancy, but differences may stem from variations in the chronicity, developmental timing, the extent of the deprivation, and brain regions studied. For instance, LB involves a chronic and multifaceted form of deprivation of “expected experiences”^11^, including erratic maternal care, poor growth, and limited nesting and bedding materials, beginning at birth (P0) and extending through P25^28,29,38^. In contrast, monocular deprivation typically lasts only four days (P28–P32) and affects visual input from just one eye^46^. Synaptic connectivity and pruning in the visual cortex are nearly mature by P28, whereas these processes remain relatively immature in the hippocampus during the second postnatal week^29,38,46^. Moreover, the increase in microglial ramification seen with monocular deprivation is both rapid (within 12 hours) and transient, disappearing by the fourth day^46^. Thus, the microglial response may be better explained by an acute shift in neuronal activity within a cortical region primed for binocular competition, rather than by deprivation per se.

We previously showed that limited bedding (LB) reduces expression of the microglial receptor TREM2, and that decreasing TREM2 levels leads to a dose-dependent decline in both in vivo and ex vivo microglial phagocytic activity in normally developing mice^38^. These findings align with other studies showing that TREM2 is exclusively expressed by microglia and is essential for normal synaptic pruning in the developing hippocampus^35,36,48^. TREM2 is also necessary to support normal energetic demands in CA1 pyramidal neurons providing yet another mechanism for impacting hippocampal function^49^. However, the specific role of TREM2 in mediating microglial function and the synaptic and hippocampal deficits observed in adolescent LB males has not been fully elucidated.

Here, we demonstrate that microglial cell volume, phagosome size, and the number of PSD95 puncta engulfed by microglia were similarly reduced in Trem2-WT and Trem2-KO mice exposed to LB, suggesting that TREM2 activity is largely absent in LB-exposed microglia—a finding corroborated by our immunohistochemical staining (Figs. 1 and S1). Moreover, LB impairs microglial volume and phagocytic activity via both TREM2-dependent and TREM2-independent mechanisms. Specifically, LB reduced microglial volume and PSD95 engulfment in Trem2-WT mice by approximately 60%, with roughly half of this reduction attributable to TREM2 loss under control conditions (i.e., TREM2-dependent) and the other half observed in Trem2-KO mice exposed to LB (i.e., TREM2-independent). Thus, LB impairs microglial synaptic pruning through additional, TREM2-independent pathways. TREM2-dependent mechanisms are relatively well characterized and involve recognition and engulfment of non-functional synapses tagged with "eat-me" signals such as phosphatidylserines. TREM2 activation also triggers intracellular signaling cascades that enhance glucose and ATP metabolism, cytoskeletal remodeling, cell motility, and the expression of pro-phagocytic genes—collectively increasing phagocytic efficiency^50–52^. In contrast, the mechanisms underlying TREM2-independent effects remain unclear, though likely candidates include P2RY12^53^, MERTK^54,55^, CD11b^38^, and SIRPα^56^.

Interestingly, TREM2 appears to play a more prominent role in regulating phagosome size. Approximately 50% of the LB-induced reduction in phagosome volume was TREM2-dependent, whereas only ∼8% was attributable to TREM2-independent mechanisms. The developmental pathways through which TREM2 influences phagosome size and function remain to be fully defined. One possibility involves direct internalization and trafficking of TREM2 itself into the phagosome. Alternatively, TREM2 may regulate phagosome maturation via activation of the PI3K/AKT/mTOR pathway^50^, as mTOR signaling enhances the expression of genes involved in phagosome maturation^57^.

Despite some microglial impairments being TREM2-independent, overexpression of TREM2 was sufficient to normalize TREM2 protein levels, microglial cell volume, the number of PSD95 puncta engulfed by microglia, and phagosome size in P17 LB-exposed pups. TREM2 overexpression also increased the body weight of LB mice to levels comparable to normally developing controls. This finding is consistent with prior work showing that chemogenetic activation of microglia during the second postnatal week similarly increased body weight in LB pups^29^, suggesting that impaired synaptic pruning— perhaps within the hypothalamus—may contribute to the growth deficits observed under LB conditions.

TREM2 overexpression further ameliorated several hippocampus-related abnormalities in adolescent LB male mice, including deficits in contextual fear conditioning, the synaptic maturity index, and glutamatergic synapse density. Using unbiased resting-state fMRI, we replicated earlier findings of reduced local functional connectivity in the dorsal hippocampus of LB mice—a reduction that was rescued in mice overexpressing TREM2. Notably, the LB-induced decrease in connectivity was more pronounced in the right hippocampus, mirroring lateralized effects observed in the synaptic maturity index. A similar paradigm causes a selective volumetric reduction in the left—but not the right—hippocampus in rats^58^, and a growing body of literature highlights important structural and functional differences between the left and right hippocampi^59^. TREM2 overexpression had no detectable impact on contextual fear conditioning or hippocampal connectivity in female littermates, consistent with previous findings that females are less susceptible to disruptions in microglial-mediated synaptic pruning^29^. LB also increased dendritic varicosities in CA1 apical dendrites of male mice, a structural abnormality that was reversed by TREM2 overexpression. Future studies are needed to determine whether these varicosities reflect underlying deficits in mitochondrial mass or energy metabolism, as has been reported in both Trem2 knockout and heterozygous mice^49^. Together, these findings identify microglial TREM2 as a key regulator of synaptic connectivity and cognitive outcomes in a mouse model of childhood neglect and deprivation.

This is the first study to demonstrate that postnatal enrichment enhances microglial-mediated synaptic pruning in the developing hippocampus of LB-exposed mice—a process that depends on the upregulation of the TREM2 receptor. These findings suggest that TREM2 plays a critical role in mediating the interaction between neuronal activity and microglial recruitment and remodeling during a key developmental window. More broadly, they reveal a novel function for microglial TREM2 in regulating synaptic pruning in response to early-life deprivation and subsequent enrichment.

## Methods Animals

All procedures involving animals were approved by the Yale University Institutional Animal Care and Use Committee (IACUC) and adhered to NIH guidelines for the care and use of laboratory animals. Studies were initially conducted using BALB/cByj (BALB/c) mice (Stock #001026, Jackson Laboratories) and then replicated using C57BL/6J mice (C57BL, Jackson Laboratories stock #000664). We follow the guidelines from the International Committee on Standardized Genetic Nomenclature for Mice^60^ in which gene and mRNA names are Italicized and the first letter capitalized (*Trem2*) versus protein name which is capitalized throughout and not italicized (TREM2). *Trem2* knockout mice (Stock # 027197, Jackson Laboratories) and *Trem2* overexpressing mice^39^ (*Trem2-OE*, Stock # 031880, Jackson Laboratories) were maintained on a C57BL/6J background. Heterozygous *Trem2-OE* males (C57BL background) were crossed with BALB/c females to generate mixed litters containing *Trem2-WT* and *Trem2-OE* offspring. These litters were then used to assess the effects of rearing conditions and Trem2 overexpression on microglia-mediated synaptic pruning at P17, as well as on contextual fear conditioning and synaptic connectivity in adolescent mice at P33–P35. Mice were housed in a temperature- and humidity-controlled vivarium (23 ± 1°C, 43% ± 2%) on a standard 12:12 hour light/dark cycle (lights on at 7:00 AM), with ad libitum access to food and water.

## Postnatal Rearing Conditions

Postnatal rearing conditions followed previously published protocols^29,38,61^. In brief, pregnant dams were housed in maternity cages containing 2 cups of corncob bedding, no nesting material, and 2–3 chow pellets placed on the cage floor. On the day of birth (P0), litters were culled to 5–8 pups and randomly assigned to one of three rearing conditions: control (CTL), limited bedding (LB), or limited bedding plus toys (LBT). CTL litters received 500 cc of corncob bedding, 15 cc of soiled bedding from the birth cage, and one 5 × 5 cm cotton nestlet from birth or postnatal day 0 (P0) until P25. LB and LBT litters received 125 cc of corncob bedding, 15 cc of soiled bedding, and no nestlet. LBT litters were provided with toys in the home cage from P14–P17, including a Safe Harbor (Lab Supplies), three colored marbles (Amazon), and two wooden blocks (Petco). Bedding was changed on P7, P14, and P21. Mice were weaned on P26 and housed with 2–3 same-sex littermates under standard conditions.

## Contextual fear conditioning (CFC)

CFC was tested in adolescent P33-35 mice using a Med Associates’ fear conditioning chamber as previously described^21,29^. Briefly, during the first day of training, mice were allowed to explore a fear conditioning chamber (Cat # VFC-008) equipped with a grid floor in the presence of a 0.4% lemon scent for 300 sec. After 300 sec of free exploration, mice were exposed to 5 shock-tone pairings at variable inter-trial intervals (30-180 sec). These were presented as 30-sec discontinued tones (7500 Hz, 80 dB) that co-terminated with a 1-sec 0.65 mA foot shock. Animals with less than 10% freezing on day 1 were excluded. On the second day, mice were returned to the same box, and contextual freezing was determined for 300 sec in the absence of a tone or a shock. No more than 2 mice per sex were tested from each litter, and all behavioral tests had at least 8 litters per sex and rearing group (the number of mice and litters for each study is available in the figure legends).

## Tissue collection

Tissue was collected from prepubescent P17 or adolescent P33-35 mice between 11:00-14:00 to minimize the diurnal effects of corticosterone. Tissue from adolescent mice was collected 24hrs after the second day of contextual fear conditioning. Mice were anesthetized with chloral hydrate (400 mg/kg) and transcardially perfused with ice-cold PBS/heparin (50 u/ml) solution (Bio-Rad, Cat #1610780; Sigma, Cat# H3393), followed by 10% formalin (Polyscience, Cat# 08279-20). P17 brains were postfixed with 10% formalin for 72hrs at 4°C and P33-35 brains were postfixed for 1hr at room temperature. After post fixation, brains were stored in PBS containing 0.05% azide at 4°C until they were processed for immunohistochemistry or DiOlistic labeling.

## Immunohistochemistry

Fifty-micron coronal sections were collected using a VT1000S vibratome (Leica) in 6 pools, each containing 5-6 slices spanning the dorsal hippocampus. Sections were washed in TBST buffer [1X Tris-buffer saline (Bio-Rad, cat#1706435), 0.5% Triton X-100 (American Bio, CAS 9002-93-1)] and then blocked with TBST containing 10% normal goat serum (NGS, Jackson ImmunoResearch, 005-000-121). To assess microglial phagocytic activity, one pool of slices was stained with rabbit anti-IBA1 (1:500; Wako, Cat. #019-19741), mouse anti-PSD95 (1:100; Merck-Millipore, Cat. #MAB1596), and rat anti-CD68 (1:400; Bio-Rad, Cat. # MCA1957T) overnight at 4°C in TBST-0.5% Triton X-100 with 3% NGS. TREM2 expression was determined using rabbit anti-IBA1 (1:500; Wako, Cat. #019-19741) and rat anti-TREM2 (1:500, R&DSystems, Cat# MAB17291) overnight at 4°C in TBST-0.2% Triton X-100 with 3% NGS. To assess the density of functional glutamatergic synapses, slices were incubated with guinea pig anti-Vglut2 (1:700; EMD-Millipore Cat. #AB2251-I), and mouse anti-PSD95 (1:100; Merck-Millipore, Cat. #MAB1596) overnight at 4°C in TBST-0.5% Triton X-100 with 3% NGS. Sections were then washed with TBST and incubated with the appropriate AlexaFluor secondary antibodies (1:400; Invitrogen) for 1 hour in the dark. Sections were washed again with TBST, mounted with Vectashield antifade (Vector Laboratories, Cat#10955), and coverslipped.

## DiOlistic labeling

This procedure was performed as described previously^29,62,63^. In brief, perfused brains were coronally sectioned at 180 µm using a VT1000S vibratome (Leica). Sections containing the dorsal hippocampus were labeled with a Helios gene gun (Bio-Rad) using 200 psi and 1.1 µm tungsten particles coated with fluorescent 1,1’-Dioctadecyl-3,3,3’,3’-Tetramethylindocarbocyanine Perchlorate (DiI, Thermo Fisher Scientific; D-282). The DiI was allowed to diffuse throughout the slices at 4°C overnight and then washed with PBS. The slices were then postfixed with 10% formalin for 30 min at room temperature, washed with PBS, mounted on Superfrost Plus slides (Thermo Scientific, Cat. # 4951F-001) with Fluoromount-GTM solution (Invitrogen, Cat. # 00-4958-02), and coverslipped.

## Resting state fMRI (rsfMRI)

Local functional connectivity maps were determined as described previously^29^. Mice were initially anesthetized with 2-3 % isoflurane, and a PE 50 tubing was placed into the i.p. cavity for dexmedetomidine (sedative) infusion. Thereafter, mice were maintained under complete anesthesia with 0.25% isoflurane and 250 μg/kg/h i.p., of dexmedetomidine. The choice of anesthetic agent was based on preliminary studies indicating that this approach provides a stable and reproducible connectivity map, and previous work has shown good agreement with outcomes observed in awake rodents ^64,65^. Body temperature was maintained at 36°C–37°C using a heating pad and monitored using an MRI-compatible rectal probe throughout the MRI experiments. The respiration rate was continuously monitored (SA Instruments Inc., Stony Brook, NY, USA) throughout the MRI experiments. Dynamic blood oxygenation dependent (BOLD) data were obtained using a Bruker 9.4T/16 magnet (Bruker BioSpin, MA, USA) using a single-shot gradient echo, echo planar imaging (GE-EPI) sequence with the following parameters: TR of 1000 ms, TE of 12 ms, in-plane resolution of 400 × 400 μm and slice thickness of 1000 μm, for a total of 300 images for each run. BOLD time series across all voxels were detrended using a second order fit and bandpass filtered (0.001 to 0.1 Hz) to exclude slow drift of the signal. Images were then registered to a brain template (200 x 200 x 200 um spatial resolution) followed by spatial Gaussian filtering (FWHM=1.5 mm) as previously described^29^.

## Microscopy and image analysis

Microscopy and image analyses were conducted as previously described^29^. To assess microglial cell volume and phagocytic activity in vivo, Z-stack images from the dorsal stratum radiatum region of the hippocampus were collected with an Olympus FV-3000 microscope with a 60X objective, 2x digital zoom and 0.30 μm intervals for a total thickness of 15-20 μm with 2x line averaging and sequential scanning. The acquired images were deconvoluted and processed using the Imaris version 9.9.1 (Oxford Instruments) according to the following protocol. A ROI (45 µm x 45 µm x10 µm) was selected using the cropped function to create 3D reconstruction surfaces in each channel using a semi-automated threshold, fixed voxel, and automated background subtraction. PSD95 puncta were detected and measured using the spot function with a threshold of 0.3 µm. IBA-1 and CD68 volumes were calculated using the surface function in Imaris. The number PSD95 puncta, CD68 phagosome volume, and TREM2 volume within Iba1+ cells were determined using the ‘spot closed to surfaces’ function with a threshold set at zero. This protocol ensures that all these cellular targets are fully embedded inside microglia. Iba1 volume, TREM2 and CD68 volumes inside microglia, as well as the number of PSD95 puncta inside microglia and inside the CD68 phagosome were averaged across 5-6 microglia cells examined per animal, and these averages were used for the analysis.

Spine density and morphology were assessed in the secondary/tertiary apical dendrites from CA1-stratum radiatum of the dorsal hippocampus (right and left hemispheres) and imaged with a Zeiss LSM 880 confocal workstation equipped with Airyscan using 63X objective at 0.46 μm intervals^29^. Dendrites that were less than 1.0 μm in diameter were then cropped to 30 μm ± 0.20 μm in average length and modeled using the semi-automated filament tracer tool in Imaris.

For each dendrite, its maximum and minimum diameter were measured using the 2D Slice tab and the average diameter was calculated and introduced in the tracer tool to draw the filament. The thresholds used for spines to be quantified were: 0.15 μm for thinnest spine head and 3.0 μm for maximum height. The default settings of the Classify Spines X Tension tool in Imaris were subsequently used to classify spines as mushroom, long thin, or filopodia. For stubby spines, the default formula was modified to 0.5 μm < length(spine) < 1.0 μm. To calculate the total spine density all spine categories were summed and divided by the length of the dendrite. Maturity index was calculated as the density of mushroom spines divided by all other spine categories. The data from 6-8 dendrites from 4 different hippocampi sections (2 left and 2 right hippocampi) were averaged to obtain the total spine density and maturity index for each individual animal.

Glutamatergic synapse density in the stratum radiatum was determined by first obtaining Z-stacks confocal images with an Olympus FV-3000 microscope, 60x objective, 2x zoom, at 0.30 µm intervals, totaling 15-20 µm in thickness. Images were processed in Imaris 9.9.1, using a 25 µm × 50 µm × 10 µm region of interest, and functional synapses defined as pre- and post-synaptic puncta located within 0-250 µm of each other. Results were exported to Excel and averaged across 6-8 fields per brain.

To calculate the percentage of hippocampal tissue showing significant differences in local functional connectivity, the local connectivity maps from three slices were overlaid with hippocampal borders using a registered brain template (Fig. 6C). The percent overlap was then calculated for each hemisphere and slice using Imaris (Fig. 6D).

## Statistical analysis

The data were carefully screened for inaccuracies, outliers, normality, and homogeneity of variance using SPSS (IBM Corp. version 26) and visualized with GraphPad Prism (iOS, version 10.0). Since no significant effect of sex or interaction were observed for microglial function, we used similar number of males and females in all studies examining the impact of rearing on microglial function and TREM2 expression. Therefore, Student -t-tests were used to confirm the effects of LB on microglial phagocytic activity and TREM2 expression in C57BL/6J 17-day-old pups (Fig 1). Two-way ANOVA was used to assess the effects of rearing (CTL vs. LB), genotype (e.g. *Trem2-KO* or *Trem2-OE*), and their interaction on microglial volume and phagocytic activity at P17 (Figs 2-3). Significant interaction between rearing and genotype was followed by Tukey-HSD post-hoc comparison across all groups. Three-way ANOVA was initially used to assess the effects of rearing (CTL vs LB), sex, and genotype (*Trem2-WT* vs. *Trem2-OE*) on contextual freezing in adolescent mice. However, since there was a significant interaction between rearing and sex, we conducted separate two-way analyses in males and females. Significant interaction was followed by Tukey-HSD post-hoc comparisons across all groups. Two-way ANOVA was also used to determine the effects of rearing (CTL vs LB) and genotype (*Trem2-WT* vs. *Trem2-OE*) on spine density, maturity index, and density of glutamatergic synapses in the stratum radiatum of adolescent male mice (Figs 5-6). One-way ANOVA was used to determine the effects of rearing (CTL vs. LB vs LBT) on microglial volume, phagocytic activity, and TREM2 expression in P17 pups, with separate analyses conducted in *Trem2-WT* and *Trem2-KO* mice (Fig 7). Local functional connectivity maps were calculated for each voxel using threshold correlation (Tc) > 0.6, a signal-to-noise ratio (TSNR> 0.5), and distance < 1 mm in individual space and normalized to a z score with the Fisher r to z transformation. Significance was assessed using Student’s t-test comparisons for different rearing conditions for the same sex and genotype (e.g. CTL-*Trem2-WT* males vs. LB-*Trem2-WT* males, Fig 6 A-B and Fig S4). Significant rearing effects were then corrected with Benjamini-Hochberg correction for multiple comparisons with a false discovery rate (FDR) < 0.05 and a local cluster size of k> 25 voxels.

## Acknowledgements

This work was supported by: NIMH R01MH136490 (AK, FH, BGS), NIMH R01MH130825 (AK), the Clinical Neuroscience Division of the VA National Center for PTSD (AK), JMM was funded by FPU21/01318 (MICIU/AEI, Spain), Plan Propio I+D+i-Movilidad (IBIMA Plataforma BIONAND, Málaga, Spain) and Acción 321 PPID (Universidad de Málaga-Banco Santander, Spain). XWY is supported by NIA RO1 AG056114, and by the Terry Semel Chair for Alzheimer’s Disease Research and Treatment from UCLA.

## Author Contributions

SA performed the immunohistochemistry and image analysis for data shown in Figs. 3, 5, and 6, wrote the first draft, and edited the final version of the manuscript. CB conducted the immunohistochemistry and image analysis for data shown in Figs. 1, 7, S1, and S5, contributed to writing the methods, and edited the final version. JMM carried out the DiOlistic studies presented in Figs. 5, 6, and S3, wrote sections of the methods, and edited the final version. SJ performed the immunohistochemistry and image analysis for data shown in Fig. 2 and collected the behavioral data presented in Figs. 4 and S2. BGS designed and conducted the rsfMRI studies shown in Figs. 6 and S4, wrote the related method sections, and edited the final version. LG and ZMK assisted with early life manipulations, tissue collection, and manuscript editing. FH contributed to the design of the imaging studies and manuscript editing. XWY generated the *Trem2-OE* mice and edited the manuscript. AK conceptualized and designed the experiments and wrote the manuscript.

## Competing Interests

The authors declare no conflict of interest.

Supplemental figures, Ahmed et al., 2025

---

**Figure S1.**
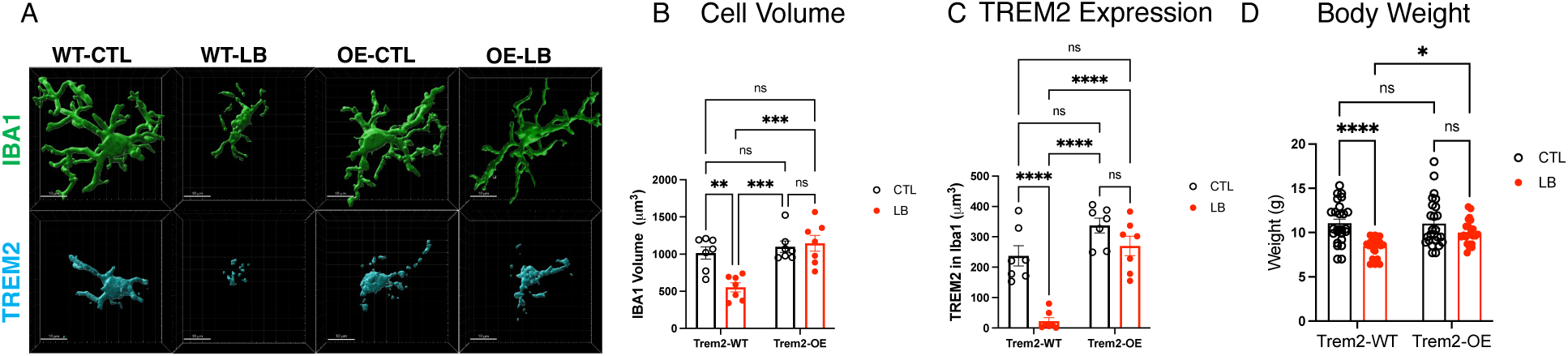
TREM2 Overexpression Normalizes Microglial Cell Volume, TREM2 Levels, and Body Weight in P17 LB Pups. Trem2-WT and Trem2-OE littermates were exposed to either control (CTL) or LB conditions and perfused at P17 to assess microglial cell volume, TREM2 expression, and body weight. (**A**) Representative Imaris images showing Iba1 (green) and TREM2 (cyan) labeling. (**B**) Cell volume: Rearing × genotype interaction: F(1, 24) = 9.274, P = 0.0056. (**C**) TREM2 expression: Interaction: F(1, 24) = 7.658, P = 0.011. (**D**) Body weight: Interaction: F(1, 93) = 3.86, P = 0.052. Data are mean ± SEM and analyzed using 2-way ANOVA followed by Tukey-HSD post-hoc tests. Significance: NS (P > 0.05), P < 0.05 (*), P < 0.01 (**), P < 0.001 (***), P < 0.0005 (****). (B–C) N = 7 mice/group from 3–4 litters (balanced for sex). (D) N = 23–25 mice/group from 10–12 litters (balanced for sex).

**Figure S2.**
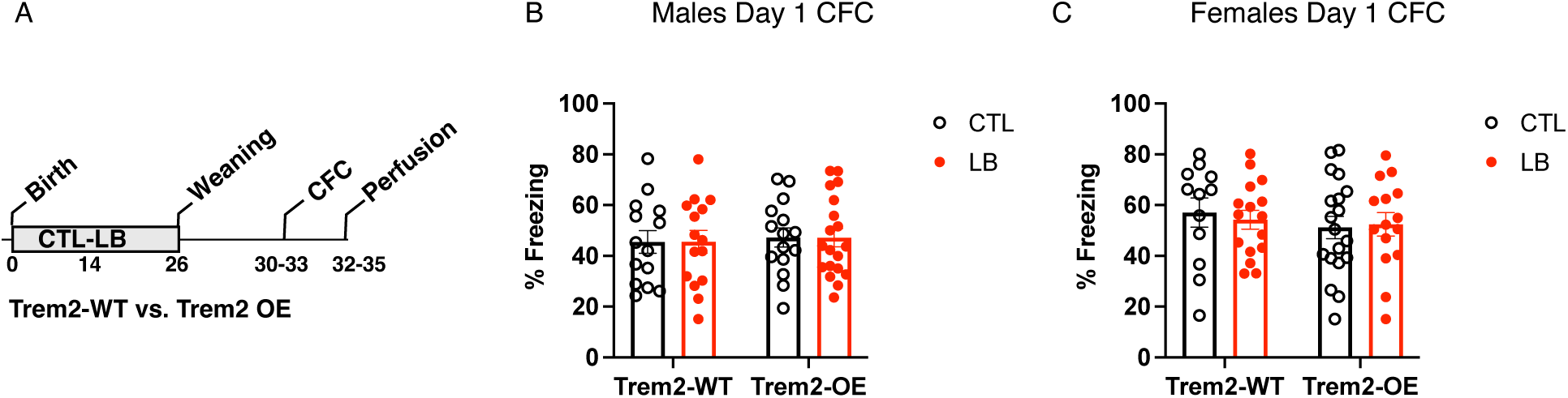
Effects of Rearing and TREM2 overexpression on Freezing Behavior During Day 1 of Training. **(A)** Experimental timeline. Freezing behavior during day 1 training in the contextual fear conditioning (CFC) in adolescent males (B) and adolescent females (C). Three-way ANOVA: sex: F (1, 117) = 5.75, P = 0.018, all other NS (genotype: F (1, 117) = 0.12, P = 0.72, rearing: F (1, 117) = 0.015, P = 0.90, sex x genotype: F (1, 117) = 0.81, P = 0.36, sex x rearing: F (1, 117) = 0.015, P = 0.90, genotype x rearing: F (1, 117) = 0.10, P = 0.74, sex x genotype x rearing: F (1, 117) = 0.11, P = 0.73. (B) Males: two-way ANOVA, Rearing: F (1, 59) = 8.84 e^-007^, P = 0.99, Genotype: F (1, 59) = 0.17, P = 0.68, Interaction: F (1, 59) = 9.124 e^-005^, P = 0.99. (C) Females: two-way ANOVA, Rearing: F (1, 58) = 0.027 P = 0.86, Genotype: F (1, 58) = 0.69, P = 0.41, Interaction: F (1, 58) = 0.19, P = 0.66. N = 12–19 per rearing, genotype, and sex condition.

**Figure S3.**
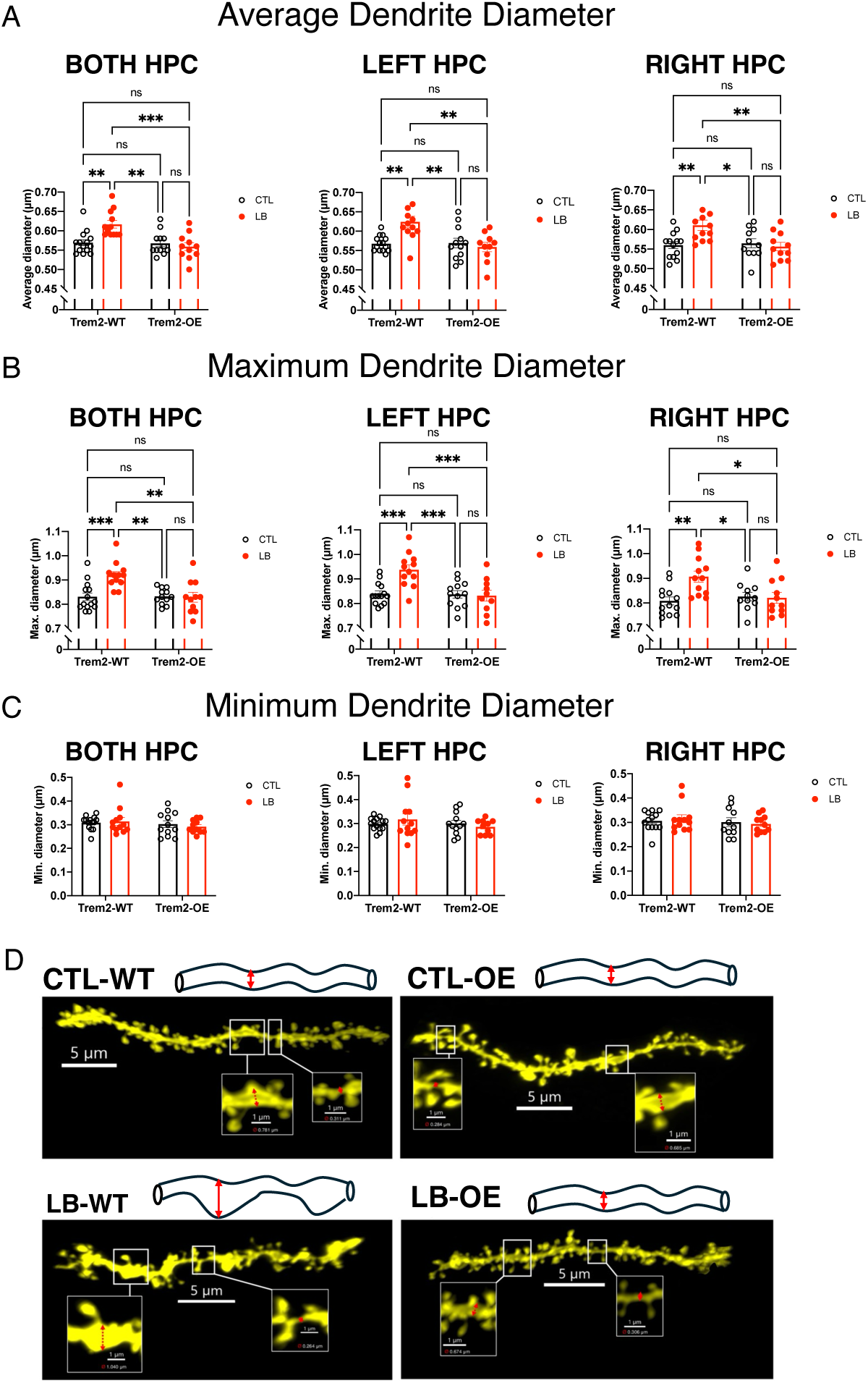
TREM2 Overexpression Normalizes Dendritic Morphology in the Left and Right Hippocampi of Adolescent LB Male Mice. Trem2-WT and Trem2-OE male littermates were exposed to CTL or LB conditions from P0–P25 and analyzed using DiOlistic labeling at P33–P35. (A-C) Diameters of apical secondary/tertiary dendrites in the dorsal CA1 stratum radiatum: (**A**) Average dendrite diameter: Both hippocampi: Rearing × genotype interaction: F(1, 45) = 9.53, P = 0.0035; Left: F(1, 44) = 8.81, P = 0.0048; Right: F(1, 43) = 6.99, P = 0.011.(**B**) Maximum diameter: Both: F(1, 45) = 8.63, P = 0.0052; Left: F(1, 44) = 9.05, P = 0.0043; Right: F(1, 43) = 7.15, P = 0.011. (**C**) Minimum diameter: No significant main effects or interactions in either hemisphere. (**D**) Representative DiOlistic images and schematics of apical dendrites. Data are mean ± SEM and analyzed by 2-way ANOVA with Tukey-HSD post hoc tests. Significance: NS (P > 0.05), P < 0.05 (*), P < 0.01 (**), P < 0.001 (***), P < 0.0005 (****). (A–C) N = 11–14 mice/group, all males.

**Figure S4.**
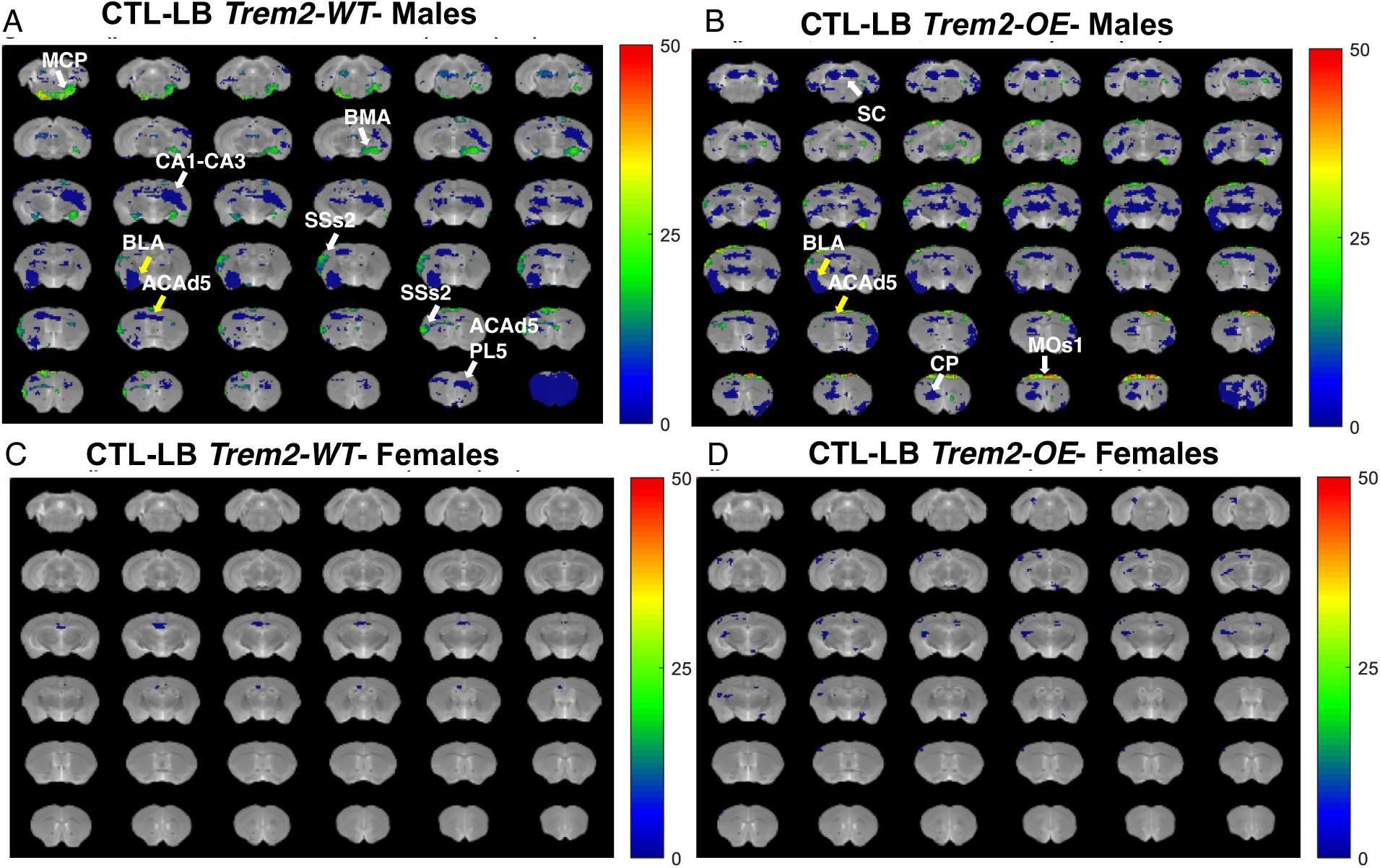
**Effects of Rearing, Sex, and TREM2 Overexpression on Local Functional Connectivity**. Trem2-WT and Trem2-OE male and female littermates were raised under CTL or LB conditions from P0–P25 and analyzed using resting-state fMRI at P33–P35. Blue and green colors indicate brain regions with significant reductions in local functional connectivity (P < 0.05, FDR corrected, local cluster size k > 25 voxels). (**A**) CTL vs LB Trem2-WT males: white arrows indicate brain regions with reduced local connectivity in LB Trem2-WT mice that were not observed in Trem2-OE males: ACAd5/PL5, SSs2, BMA, MCP, CA1-CA3; yellow arrows show brain regions reduced in LB regardless of genotype: BLA, ACAd5. (**B**) CTL vs LB Trem2-OE males: white arrows indicate regions reduced in LB Trem2-OE but not Trem2-WT: MOs1, CP, SC; yellow arrows show regions reduced regardless of genotype: BLA, ACAd5. (**C**) CTL vs LB Trem2-WT females: no significant changes. (**D**) CTL vs LB Trem2-OE females: no significant changes. A-B are the same data shown in Fig. 6 A-B. N = 6 mice per sex, genotype, and rearing group. Abbreviations: ACAd5: Anterior cingulate area; PL5: Prelimbic area; SSs2: Supplemental somatosensory area; BLA: Basolateral amygdalar Nucleus; BMA: Basomedial amygdalar Nucleus; MCP: Middle cerebellar penducle; CA1-CA3: Hippocampal area; MOs1: Secondary motor area; CP: Caudoputamen; SC: Superior colliculus.

**Figure S5.**
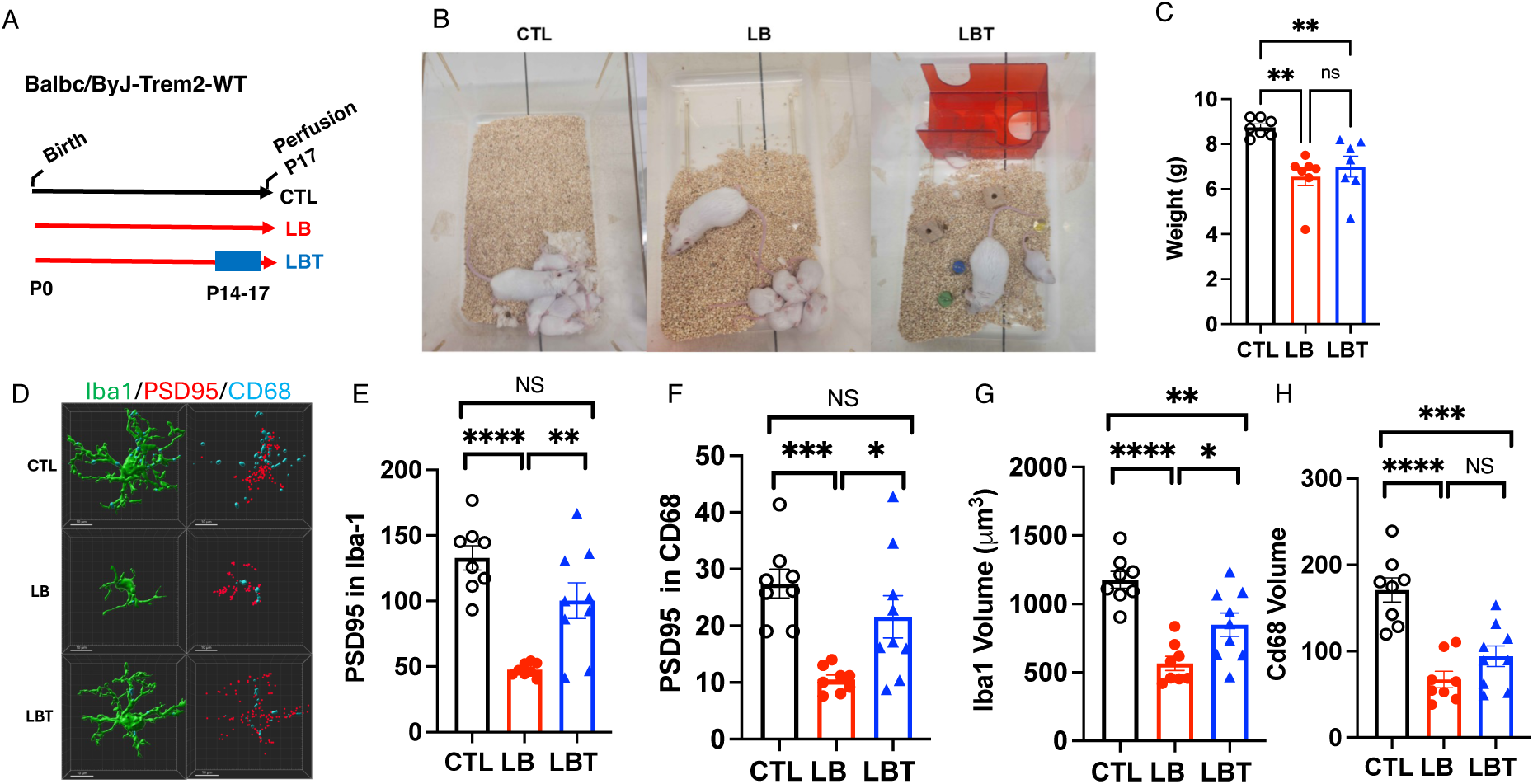
Brief Postnatal Enrichment Enhances Microglial-Mediated Synaptic Pruning in P17 BALB/cByJ Pups. BALB/cByJ litters were randomized to CTL or LB conditions. At P14, some LB cages received toys (LBT), and all mice were perfused at P17 to assess body weight and in vivo phagocytic activity of microglia in the stratum radiatum. (**A–B**) Experimental timeline and rearing conditions. (**C**) Body weight: F (2,18) = 9.75, P = 0.0014.(**D**) Representative Imaris reconstructions of microglia stained with Iba1 (green), PSD95 (red), and CD68 (cyan). (**E**) PSD95 puncta within microglia: F (2,22) = 17.89, P < 0.0001.(**F**) PSD95 puncta within phagosomes: F (2, 22) = 9.29, P = 0.0012. (**G**) Microglial volume: F (2,22) = 18.69, P < 0.0001. (**H**) Phagosome volume: F (2,22) = 19.39, P < 0.0001. Data are mean ± SEM and analyzed using one-way ANOVA followed by Tukey-HSD post-hoc tests. Significance: NS (P > 0.05), P < 0.05 (*), P < 0.01 (**), P < 0.001 (***), P < 0.0005 (****). N = 8 mice per rearing group, balanced for sex.

